# Genomic prediction using low density marker panels in aquaculture: performance across species, traits, and genotyping platforms

**DOI:** 10.1101/869628

**Authors:** Christina Kriaridou, Smaragda Tsairidou, Ross D. Houston, Diego Robledo

**Affiliations:** The Roslin Institute and Royal (Dick) School of Veterinary Studies, University of Edinburgh, EH25 9RG Midlothian, United Kingdom

**Author notes:** Corresponding authors: Diego Robledo –, Ross D. Houston –. E-mail addresses: Christina Kriaridou –, Smaragda Tsairidou –.

**Keywords:** breeding, disease resistance, growth, GBLUP, fish, oyster, salmon

## Abstract

Genomic selection increases the rate of genetic gain in breeding programmes, which results in significant cumulative improvements in commercially important traits such as disease resistance. Genomic selection currently relies on collecting genome-wide genotype data accross a large number of individuals which requires substantial economic investment. However, global aquaculture production predominantly occurs in small and medium sized enterprises for whom this technology can be prohibitively expensive. For genomic selection to benefit these aquaculture sectors more cost-efficient genotyping is necessary. In this study the utility of low and medium density SNP panels (ranging from 100 to 9000 SNPs) to accurate predict breeding values was tested and compared in four aquaculture datasets with different characteristics (species, genome size, genotyping platform, family number and size, total population size, and target trait). A consistent pattern of genomic prediction accuracy was observed across species, with little or no reduction until SNP density was reduced below 1,000 SNPs. Below this SNP density, heritability estimates and genomic prediction accuracies tended to be lower and more variable (93 % of maximum accuracy achieved with 1,000 SNPs, 89 % with 500 SNPs, and 70% with 100 SNPs). Now that a multitude of studies have highlighted the benefits of genomic over pedigree-based prediction of breeding values in aquaculture species, the results of the current study highlight that these benefits can be achieved at lower SNP densities and at lower cost, raising the possibility of a broader application of genetic improvement in smaller and more fragmented aquaculture settings.

## BACKGROUND

Aquaculture is the fastest growing food industry worldwide (FAO2018). While capture fisheries production has stagnated since the late 90s, aquaculture production has been consistently increasing 5.8 % per year since 2001 (FAO 2018), and this trend is expected to continue in the coming years to cope with the food demands of a growing human population. Nonetheless, aquaculture is still a relatively young industry, and although technological advances have been rapidly implemented to improve production volume and efficiency for some high-value species, these are slower to reach the lower-value, high-volume species that underpin most of global production. This is typified by genetic improvement technologies, where species such as Atlantic salmon have large and well-managed breeding programmes akin to those for pigs and poultry, while most aquaculture species lag significantly behind. In part, this is due to the wide diversity of aquaculture species, with the top 20 animal species accounting for less than 80 % of the total production (FAO 2019) in contrast to terrestrial livestock, where four species are the source of > 90 % of the world meat production. In addition, the majority of aquaculture takes place in small to medium-sized farms, primarily situated in low to medium income countries. This context hinders the implementation of emerging technologies to help improve production, primarily due to their prohibitive cost.

One such technology is genomic selection, which utilises genetic markers to identify the animals with the highest breeding values to select for producing the next generation (Meuwissen et al. 2001). Selective breeding programmes are being increasingly utilised for aquculture species, and have been shown to be highly effective in improving production traits, especially growth (Gjedrem and Rye, 2018). Genomic selection consistently outperforms family-based selection based on pedigree only (Zenger et al. 2018), leading to cumulative genetic gains over generations that incrementaly enhance the performance of farmed species. One of the main reasons underlying the slow uptake of genomic selection in aquaculture is genotyping costs. Genotyping usually relies on high-density SNP array platforms, which can be prohibitively expensive for routine application for most aquaculture breeding programmes, due to the need to genotype thousands of performance tested fish (i.e. the reference population) and the selection candidates. One avenue to democratise genomic selection for smaller-scale, more fragemented aquaculture sectors is to exploit low-density SNP panels for which per-sample genotyping costs can be a fraction of the cost of SNP arrays.

However, it may be expected *a priori* that this cost-reduction due to reduced genotype density comes at the expense of reduced prediction accuracy in a breeding programme. The improved accuracy of genomic selection compared to pedigree-based approaches is primarily derived from an improved estimation of the genomic similarity between each pair of individuals. In most family-based aquaculture breeding programmes, a procedure known as sib-testing (short for sibling testing) is performed, whereby trait records are obtained from full siblings of the selection candidates – a process enabled by the high fecundity of aquaculture species. With pedigree-based selection, the genomic similarity between full-sibs is assumed to be 50 %, but the reality is that it can vary substantially around this value as a consequence of Mendelian sampling and linkage disequilibrium (Hill and Weir, 2011). In theory, the accuracy of estimating this genomic similarity should decrease as the density of genetic markers employed reduces, which would have a negative impact on prediction accuracy and consequently on genetic gain. However, in emperical studies of aquaculture species to date this decrease in accuracy seems to be relatively small and only observable once SNP densities drop to a few hundred markers (e.g. Tsai et al. 2016; Correa et al. 2017; Robledo et al. 2018; Yoshida et al. 2018; Vallejo et al. 2018; Gutierrez et al. 2018; Palaiokostas et al. 2019; Tsairidou et al. 2019), which is likely a consequence of the large full sibling family sizes, such that long haplotypes are shared between many individuals in the reference and test population.

Therefore, low density genotyping appears to be a promising solution for enabling access to the benefits of genomic selection to a broader range of aquaculture species and sectors. However, the optimal SNP density to use is unclear, and may be expected to vary depending on the species, population history and trait of interest. The goal of this study was to assess if those variables affect the performance of low-density SNP panels, and to determine if an optimal genotyping density can be identified as a practical, broad recommendation for aquaculture breeding programmes. To do so, the performance of SNP panels of varying densities in estimating genetic parameters and breeding values was tested using previously published datasets for diverse aquaculture species, phenotyped for different traits and genotyped with different platforms.

## MATERIALS AND METHODS

### Datasets and phenotypes

Genotypes and phenotypes were obtained from four previously published studies in four different species, briefly: i) Atlantic salmon (*Salmo salar*) challenged with amoebic gill disease (AGD) were phenotyped for mean gill score (subjective 0 – 5 scoring system, commonly used as a measure of gill damage) and amoebic load (real-time PCR), and genotyped using a combined salmon-trout 17K SNP array (Robledo et al. 2018); ii) Common carp (*Cyprinus carpio*) were measured for growth traits (standard length and weight), and genotyped using RAD sequencing for ~12K SNPs (Palaiokostas et al. 2018); iii) Sea bream (*Sparus aurata*) challenged with *Photobacterium damselae* (causative agent of pasteurellosis) were measured for time to death, and genotyped using 2b-RAD sequencing for ~12K SNPs (Palaiokostas et al. 2016); and iv) Pacific oyster (*Crassostrea gigas*) challenged with ostreid herpesvirus (OsHV-1-μvar) were measured for time to death, and genotyped using a SNP array with ~27K informative Pacific oyster SNPs (Gutiérrez et al. 2019).

### Quality control and low density SNP panel design

Genotypes from the four datasets were filtered with PLINK v.1.9 (Purcell et al. 2007), excluding individuals with > 20 % missing genotypes, and SNPs with > 10 % missing genotypes, deviating significantly from Hardy-Weinberg (p-value < 10^−6^) and with minor allele frequencies < 0.05. A summary of the genetic marker and trait data used for the four different datasets used in this study after quality control is shown in Table 1.

**Table 1.** Summary of the datasets.

| Species | Individuals before and after QC |  | SNPs before and after QC |  | Full-sib families | Phenotypes |
| --- | --- | --- | --- | --- | --- | --- |
| *Cyprinus carpio* | 1,214 | 1,211 | 12,311 | 6,966 | 195 | Log length, weight |
| *Crassostrea gigas* | 718 | 718 | 21,338 | 14,028 | 23 | Days to death |
| *Salmo salar* | 1,481 | 1,481 | 16,582 | 9,866 | 85 | Gill score, amoebic load |
| *Sparus aurata* | 777 | 741 | 12,085 | 7,598 | 73 | Days to death |

SNP panels of varying densities were tested by taking subsets of the full QC-filtered SNP panel for each dataset. Panels of the following densities were tested in every species: 100, 200, 300, 400, 500, 600, 700, 800, 900, 1000, 1200, 1400, 1600, 1800, 2000, 2250, 2500, 2750, 3000, 3500, 4000, 4500 and 5000. Additionally, 6,000, 7,000 and 9,000 SNP panels were tested depending on the total number of SNPs remaining after quality control (carp 6,000 SNPs; sea bream 7,000 SNPs; salmon and oyster 7,000 and 9,000 SNPs). The SNPs for each panel were selected using two different strategies (R package CVrepGPAcalc v1.0, https://github.com/SmaragdaT/CVrep/): i) random selection of SNPs within each chromosome (or linkage group for sea bream and oyster), where the number of SNPs selected from each chromosome / linkage group was proportional to its length; and ii) random selection of SNPs across the genome, where SNPs were randomly chosen irrespective of their genomic position. For each SNP density, five different SNP panels were selected to account for potential bias arising from SNP sub-set selection.

### Estimation of genetic parameters

Heritabilities of the measured traits in each dataset were estimated using ASReml 3.0 (Gilmour et al. 2014) fitting the following linear mixed model:

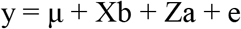

where y is a vector of observed phenotypes, μ is the overall mean of phenotype records, b is the vector of fixed effects, a is a vector of additive genetic effects distributed as ~N(0,**G**σ^2^_a_) where σ^2^_a_ is the additive (genetic) variance and **G** is the genomic relationship matrix. **X** and **Z** are the corresponding incidence matrices for fixed and additive effects, respectively, and e is a vector of residuals. The identity-by-state genomic relationship matrix (**G**) was calculated using the GenABEL R package (“gkins” function; Aulchenko et al. 2007) kinship matrix (Amin et al., 2007), multiplied by two and inverted.

The different fixed effects included in the model for each species were i) tank (2 levels) in Atlantic salmon, ii) factorial-cross group (4 levels) in carp, iii) none in sea bream, and iv) tank (2 levels) in oyster.

### Genomic prediction

The accuracy of genomic prediction was estimated by ten replicates of fivefold crossvalidation analysis (training set 80 %, validation set 20 %; R package CVrepGPAcalc v1.0, https://github.com/SmaragdaT/CVrep). The phenotypes recorded in the validation population were masked, and genomic best linear unbiased prediction (GBLUP) was applied to predict the breeding values of the validation sets in ASReml 3.0, using the linear mixed model described above. Prediction accuracy was calculated as the correlation between the predicted EBVs of the validation set and the actual phenotypes divided by the square root of the heritability estimated from the full dataset 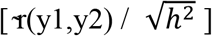.

## RESULTS

### Trait summary

In total six traits were studied. Two traits related to Atlantic salmon resistance to AGD were used, gill score (subjective values 0 – 5) and amoebic load (qPCR, Ct values), with means of 2.79 ± 0.85 and 31.36 ± 3.24, respectively. The estimated genomic heritabilities values were moderate for both phenotypes, 0.22 (± 0.04) for gill score and 0.24 (± 0.04) for amoebic load. Two growth traits were studied in carp, length and body weight, with means of 77.01 ± 7.11 mm and 16.33 ± 4.58 g respectively. Length showed a skewed distribution, deviating significantly from normality, and therefore was log-transformed. The heritability estimates were 0.27 (± 0.04) for log-transformed length, and 0.19 (± 0.04) for carp weight. Days to death were measured in pasteurellosis infected sea bream. The mean and standard deviation of surviving days for sea bream was 10.40 ± 4.08, and the heritability was 0.20 (± 0.06). The same trait, days to death, was measured in oyster infected with OsHV-1-μvar. Survivors were assigned a value of 8 for the variable “days to death”. The mean for this trait was 6.76 ± 1.91 days, and the heritability 0.49 (±0.05).

### Reduced SNP panel densities decrease the precision of genomic heritability estimates

Low-density panels were designed from the full set of SNPs that passed the QC filters in each species using two different strategies: (i) randomly selected across the genome and (ii) randomly selected within chromosome. The results obtained with both selection strategies were similar, therefore only the results of the panels randomly selected across the genome are shown.

Heritabilities for the six traits were re-calculated using the reduced density SNP panels (Figure 1). In general, decreasing marker density led to progressively lower heritability estimates, however a clear downwards trend is only observed ~1000 bp onwards. The heritability estimates obtained for 100 SNPs decreased to 23 to 41 % of the values obtained for the full density panel, while for 200 SNPs the decrease was on average ~50 %.

**Figure 1.**
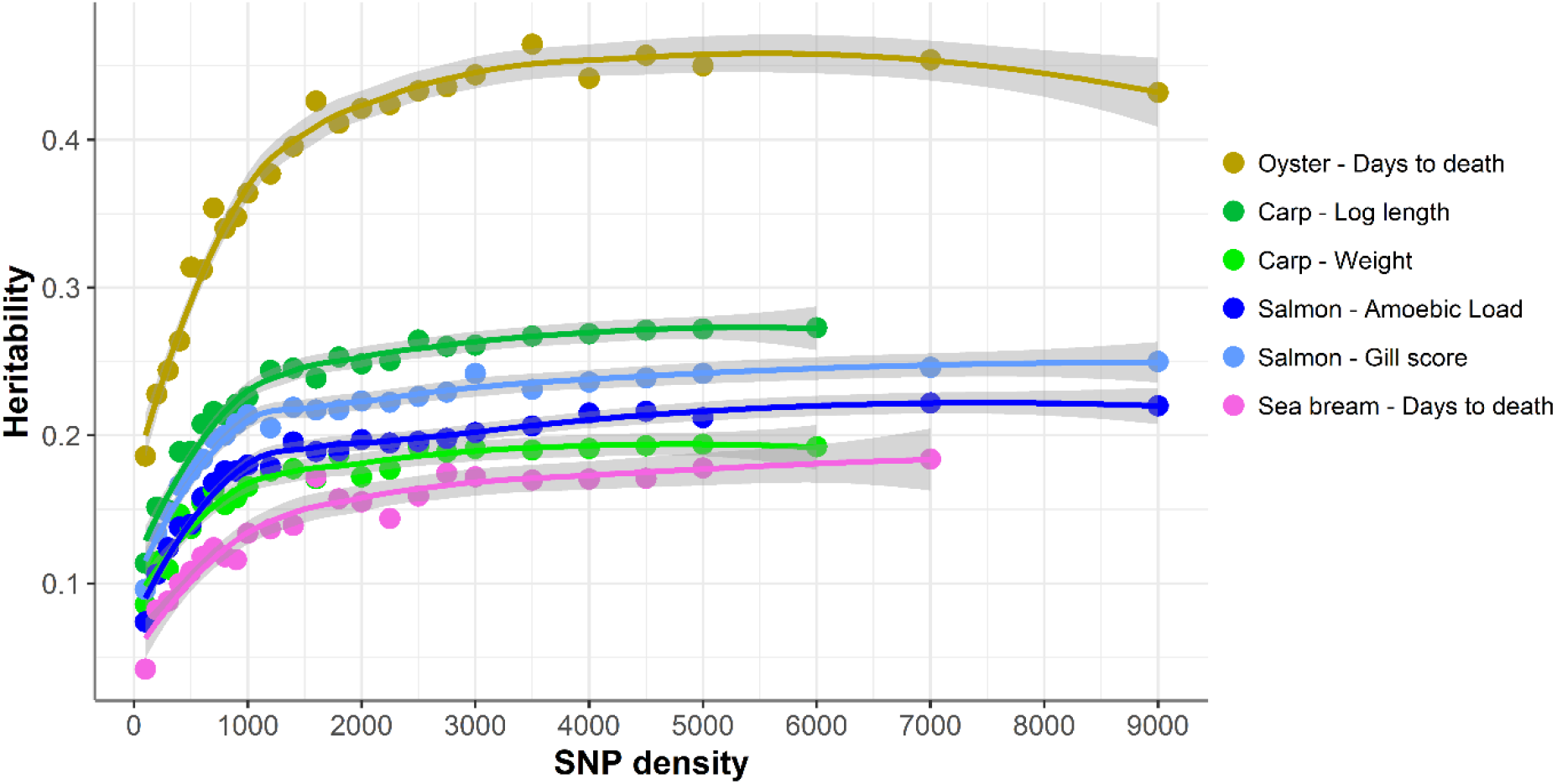
Heritability estimates using low-density panels. The heritability was calculated using a linear mixed model with the genomic relationship matrix obtained with each low-density panel. For each density we used five different low-density panels, and the average of the heritabilities of the five panels is shown. The trend line was calculated using a Loess regression (local polynomial regression, span = 0.75), and the shadow represents the confidence intervals.

### Genomic prediction using reduced SNP panels

The accuracy of genomic selection was evaluated using ten replicates of five-fold crossvalidation (training set 80%, validation set 20%) for five different panels per SNP density (Figure 2). Since the heritabilities decrease substantially with lower panel densities, the accuracy of genomic selection for all cross-validation analyses was calculated using the heritability obtained with the whole SNP panel, considered to be the most accurate heritability for the trait. Genomic selection accuracy remained practically unchanged for every dataset until marker density was reduced below ~ 2,000 SNPs, and a steep decrease was observed only for ≤1,000 SNPs. The common trend observed accross the different species, traits, and genotyping platforms is clearly observed by plotting the proportion of the full SNP panel accuracy achieved with each low-density panel (Figure 3). Despite the significant differences between datasets and traits, the genomic selection accuracies obtained with low-density panels were remarkably similar. The average proportion of the full panel accuracy achieved with 2,000 SNPs was 0.97, with 1,000 SNPs 0.93, and with 500 SNPs 0.89. With 100 SNPs the accuracy was reduced to 0.70 of that obtained with the whole density panel.

**Figure 2.**
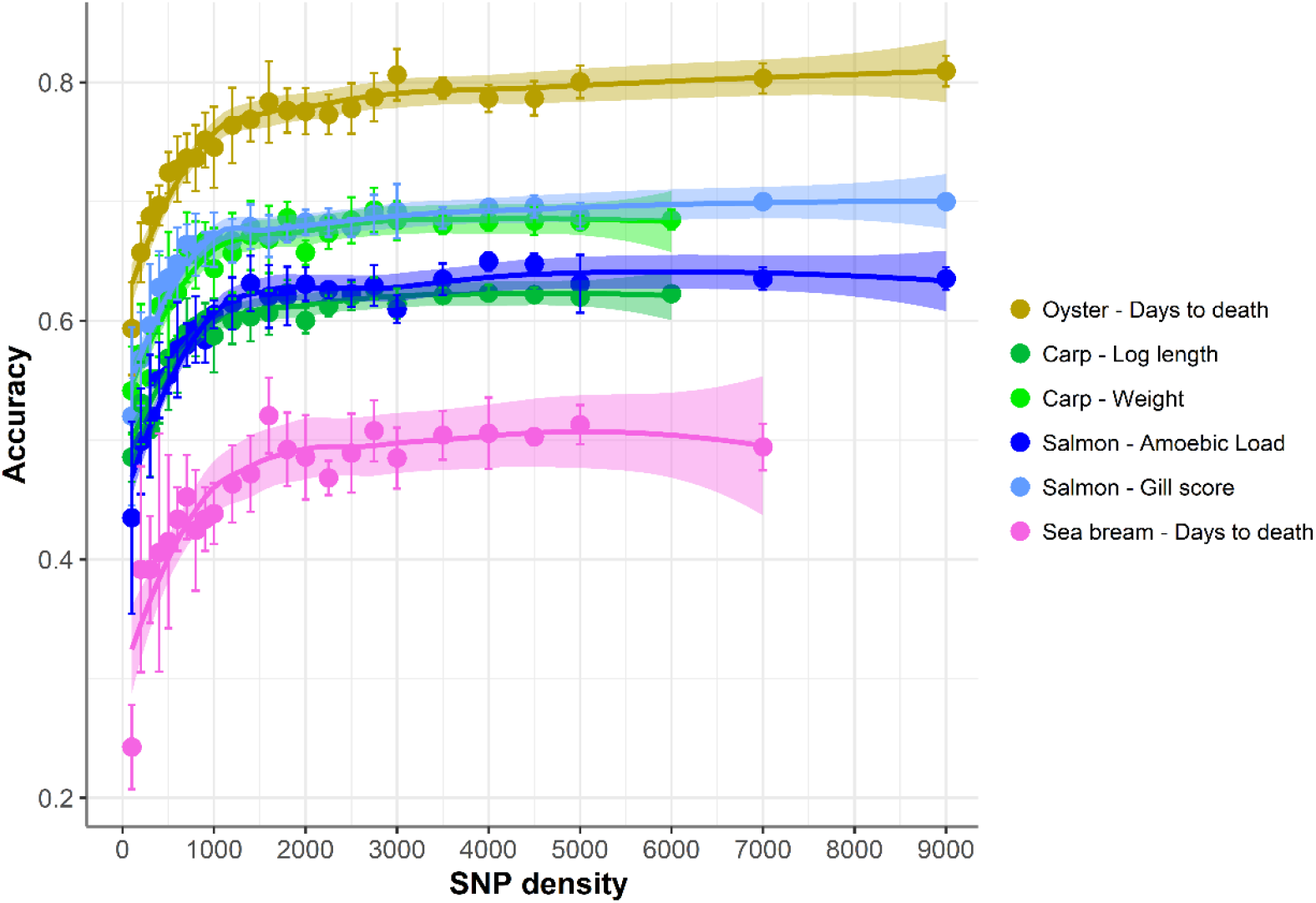
Genomic selection accuracy using low-density panels. Mean accuracy and standard deviation of genomic selection for five different SNP panels per density. The trend line was calculated using Loess regression (local polynomial regression, span = 0.75), and the shaded areas represent the confidence intervals.

**Figure 3.**
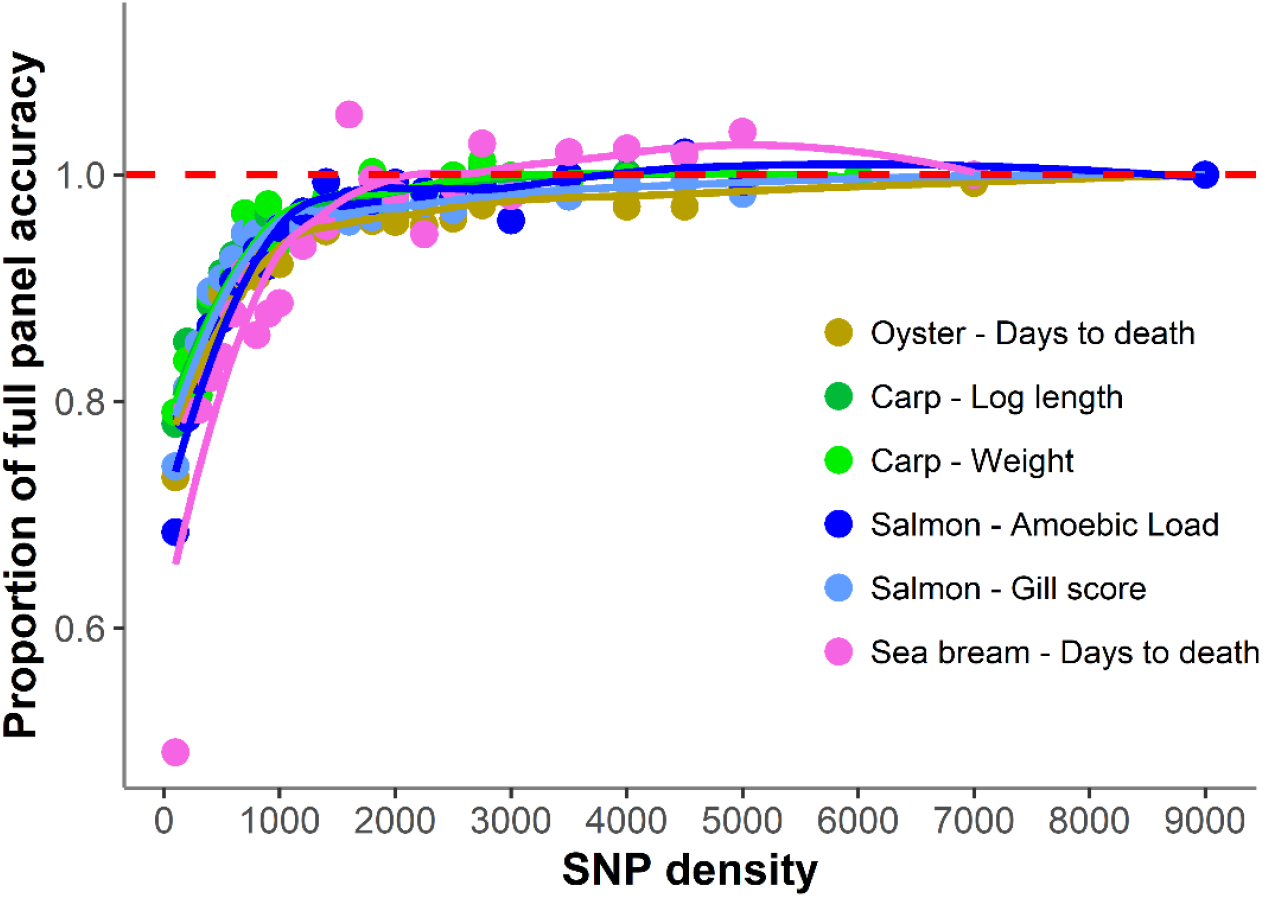
Proportion of genomic selection accuracy achieved with low-density panels. The proportion of accuracy achieved by each SNP density was calculated by dividing the mean accuracy at that density by the mean accuracy obtained using the full high density SNP panels. The trend line was calculated using a Loess regression (local polynomial regression, span = 0.75), and the shadow represents the confidence intervals.

In addition, with decreasing SNP density the differences in prediction accuracy between different replicates of SNP panels of the same density increased (Figure 4). Therefore, SNP selection seems to be more relevant for the design of low-density panels than for higher density panels. On average, the difference between the maximum and minimum accuracies achieved by 100 density SNP panels was 0.11; salmon mean gill score showed the largest difference (0.19) and carp Log standard length the lowest (0.05).

**Figure 4.**
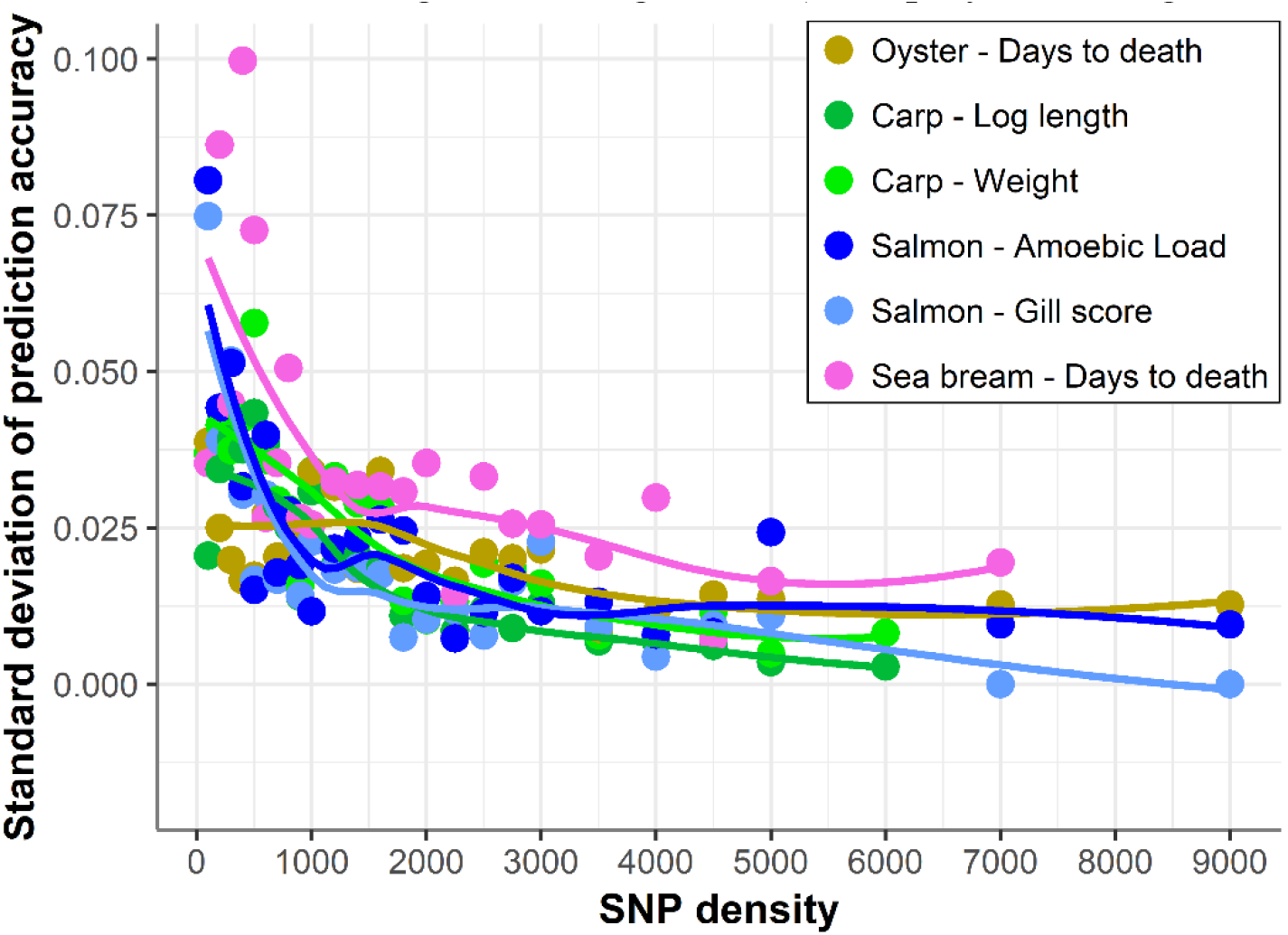
Standard deviation of selection accuracy using low-density panels. Variation in genomic selection accuracy across the different SNP panels of the same density. The trend line was calculated using a Loess regression (local polynomial regression, span = 0.75).

## DISCUSSION

Genomic selection has clear potential for improving selection accuracy and genetic gain in aquaculture breeding programmes, but the cost of genotyping can be prohibitive for many species and sectors. Therefore, since the price of per sample genotyping is generally associated with SNP density, knowledge of the lowest SNP density at which optimal genetic parameter estimation and genomic prediction can be performed is valuable. It may be expected that the optimal SNP density for genomic prediction would be species, traits, and genotyping platform-specific. In the current study, genotype and trait datasets from four diverse aquaculture species (Atlantic salmon, common carp, gilthead sea bream, and Pacific oyster), genotyped using different genotyping platforms (SNP array and RAD sequencing) were evaluated to search for common patterns of the impact of reducing SNP marker density on genomic prediction accuracy. The results were consistent across the different datasets, suggesting that a SNP panel between 1,000 and 2,000 SNPs would be sufficient for near-maximal prediction accuracy for most polygenic traits in aquaculture populations. These results and their consistency are encouraging for lower-cost genotyping, and therefore improved affordability of genomic selection across different species and aquaculture sectors.

The uniformity of the results is relatively surprising considering the notable background differences between the four datasets. The trait, genotyping platform, family structure, population size or genome size seem to be relatively unimportant factors for the performance of low density SNP panels, since genomic prediction accuracy trends were consistent across the four species. The large family sizes observed in most aquaculture species might partially explain these results. The genetic distance between training and validation populations has a large impact on the efficacy of genomic selection (accuracy decreases with increasing genetic distance; Scutari et al. 2016; Tsai et al. 2016; Tan et al. 2017; Palaiokostas et al. 2019). The underlying cause is that related individuals tend to share long haplotypes, which can be accurately captured with relatively sparse numbers of SNPs; however as genetic distance increases between training and validation populations haplotype length is reduced, and higher density panels are required to accurately capture the genomic similarity between animals. Most aquaculture species are highly fecund, and each pair of animals frequently produces thousands of offspring, meaning that inclusion of multiple full and half siblings in training and validation sets is common practice. Consequently, we consider that these results are generally applicable to polygenic traits in most aquaculture breeding schemes where close relatives of the selection candidates are routinely phenotyped.

Nonetheless, there will be situations where genomic prediction across generations or across populations is necessary. In these scenarios the shared haplotypes between pairs of individuals will be shorter, and therefore capturing genomic relatedness (if it exists; i.e. relatedness between unrelated populations will be zero and therefore of no use for prediction) is much more challenging and is likely to require higher SNP densities (Tsai et al. 2016). An avenue to increase the accuracy of low-density panels across sets of distantly related individuals could be the prioritization of variants that have a higher likelihood of directly effecting the trait in question, rather than linked markers. For example, SNPs which fall in genes or other genomic features with a direct biological effect on the trait of interest, and the utilization of selection models that exploit biological priors (MacLeod et al. 2016). However, establishing causal relations between genotypes and phenotypes is not trivial and will require extensive efforts in functional annotation of genomes (e.g. Macqueen et al. 2017), and collection of genotype and phenotype datasets across very large reference populations (Hickey 2013). Consequently, low-density panels are not likely to be a feasible option for prediction across datasets without a high degree of relationship, which would require a large number of genome-wide distributed genetic markers. Nonetheless, this scenario is rare, and in the ample majority of aquaculture breeding programmes full-sibs of the selection candidates are routinely phenotyped.

SNP panels consisting of <1,000 SNPs show a steep decline in genomic prediction accuracy, as does the estimated heritability, and the variation between replicate SNP panels of the same density increases. This suggests that low density panels are not accurately capturing the genetic relationship between animals, and that the performance of low-density SNP panels could be highly dependent on SNP choice. While leveraging additional layers of information might enable the design of high-performing low-density SNP panels, these would have to be tailored to specific breeding programmes and might require substantial investment, i.e. an initial large-scale genotyping effort and extensive time commitment to determine the best panel, or potential functional experiments to establish marker function. Further, the performance of extreme low-density panels could fluctuate across generations as allelic frequencies vary. On the contrary, genotype imputation from very low-density panels (i.e. 100-200 SNPs) to medium density (i.e. 1 K – 5 K) might be a more generally applicable strategy to achieve the optimal balance between economic cost and genetic gain. Previous studies have shown the potential of imputation to achieve near-maximal accuracies in aquaculture populations (Tsai et al. 2017; Yoshida et al. 2018); and a recent study by our group reported that imputation from 200 (offspring) to 5,000 SNPs (parents) results in selection accuracies similar to those obtained with 75K SNP panels for sea lice resistance in Atlantic salmon (Tsairidou et al. 2019). In aquaculture, studies of imputation for genomic selection have been limited to salmonid species to date, however it shows great potential and is likely to be a staple component of modern aquaculture breeding programmes.

## CONCLUSIONS

The patterns of loss of genomic prediction accuracy with reduced density SNP panels are strikingly consistent across datasets of different aquaculture species, despite their differences in population and family structure, phenotype and trait definition, and genotyping platform. These results suggest that SNP densities between 1,000 and 2,000 SNPs will frequently result in selection accuracies very similar to those obtained with high-density genotyping, irrespectively of the specifics of the breeding programme design or population structure, assuming the presense of close relatives in the training and validation sets. Further, the higher variance between SNP panel replicates observed with decreasing density suggests that nonrandom SNP selection can increase the selection accuracy of low-density panels. In summary, this study suggests that low-density SNP panels offer a cost-effective solution for broadening the impact of genomic selection in aquaculture, leading to improved enhanced performace of stocks and improved global food security.

## Data availability

All data used in this study has been previosuly published and is available in the corresponding manuscripts, namely Palaiokostas et al. 2016 (Sea bream), Palaiokostas et al. 2018 (Carp), Robledo et al. 2018 (Atlantic salmon), and Gutiérrez et al. 2019 (Oyster).

## Author’s contributions

RH and DR were responsible for the concept and design of this work. CK and ST designed and performed the genetic analyses. CK, RH and DR drafted the manuscript. All authors read and approved the final manuscript.

## Acknowledgements

The authors gratefully acknowledge funding from the Scottish Aquaculture Innovation Centre (Grant number SL_2017_09), and BBSRC Institute Strategic Programme Grants to the Roslin Institute (BB/P013732/1, BB/P013740/1, BB/P013759/1). Christina Kriaridou was supported by Erasmus+ programme of the European Union (ER/SMP-OUT/2018-0033). The European Commission’s support for the production of this publication does not constitute an endorsement of the contents, which reflect the views only of the authors, and the Commission cannot be held responsible for any use which may be made of the information contained therein. The authors also acknowledge the contribution of Alastair Hamilton and Hendrix Genetics for generation of the Atlantic salmon data; Christos Palaiokostas, Martin Kocour and Martin Prchal for generation of the carp data; Christos Palaiokostas, Serena Ferraresso, Rafaella Franch and Luca Bargelloni for generation of the sea bream data; and Alejandro P. Gutiérrez, Jane Symonds, Nick King, Konstanze Steiner and Tim P. Bean for generation of the oyster data, which was funded by Cawthron’s MBIE-funded Cultured Shellfish Programme, CAWX1315.

